# An updated genome-scale model for *Xylella fastidiosa* subsp. *pauca* De Donno

**DOI:** 10.1101/2022.11.28.518184

**Authors:** Alexandre Oliveira, Emanuel Cunha, Miguel Silva, Cristiana Faria, Oscar Dias

## Abstract

*Xylella fastidiosa* is a gram-negative phytopathogenic bacterium that caused a significant economic impact around the world. In the last decade, genome-scale metabolic models have become important systems biology tools for studying the metabolic behaviour of different pathogens and driving the discovery of novel drug targets. This work is a second iteration of the iMS508 model for *X. fastidiosa* subsp. *pauca* De Donno. The model comprises 1138 reactions, 1234 metabolites, and 509 genes. *in silico* validation of the metabolic model was achieved through the comparison of simulations with available experimental data. Aerobic metabolism was simulated properly and fastidian gum production rates predicted accurately.

## 1. Introduction

*Xylella fastidiosa* is commonly known and widespread gram-negative phytopathogenic bacterium [1]. It usually resides in the xylem of its host and is then transmitted to other plants by insects that feed on the xylem-fluid [2]. Even though host specificity is a prevalent characteristic in many phytopathogenic bacteria [3], this is not the case of *X. fastidiosa*. According to the *Xylella spp*. host plant database [4], the infection has been recorded in over 500 plant species. Still, *X.fastidiosa’s* subspecies are rather host-specific and usually related to certain plant diseases [5,6]. One key characteristic of the infection caused by this phytopathogen is that some plants may be infected whilst not showing any symptoms [6]. Hence, an *X. fastidiosa* outbreak is significantly harder to mitigate.

The *X. fastidiosa* infection was first detected in European territory in the Apulia region of Italy and was caused by the *X. fastidiosa* subsp. *pauca* De Donno [7,8]. This led to a severe outbreak of a disease that would come to be known as olive quick decline syndrome. As a result, thousands of olive trees died, thus having a negative impact on the olive oil market and the local economy [8]. Future projections indicate, in a worst-case scenario, an economic impact of 5.2 billion Euros in Italy alone for the next 50 years [9]. Hence, *X. fastidiosa* was declared a quarantine pest by the European Commission, and was the highest-ranked crop infesting pest according to the Impact Indicator for Priority Pests (I2P2) metric, leading all the economic, social, and environmental ranked domains in 2019 [10].

With the rise in popularity of high-throughput sequencing techniques, the availability of genomic and metabolic data has been exponentially increasing. As a result, fields such as Systems Biology have gained significant relevance. Systems Biology aims at studying cells through a system-level approach, trying to uncover their structure and the dynamics between each of its components [11]. Genome-Scale Metabolic (GSM) Models are Systems Biology tools that combine both genomic and metabolic data for a given organism. Throughout the years these models have been extensively used to predict phenotypes for a given organism, under specific conditions, and to drive new knowledge of the metabolic behaviour of the organism in study [12]. Moreover, GSM models are useful tools to unveil novel drug targets to kill pathogens with minimum effect on the host, leading to drug discovery or re-purposing [13,14].

In 2020, a GSM model of *X.fastidiosa* subsp. multiplex CFBP 8418 was published. That work aimed at developing a metabolic model to uncover metabolic characteristics connected to the fastidious growth of the phytopathogen [15]. Moreover, an in-house developed GSM model of *X.fastidiosa* subsp. *pauca* De Donno was published in a conference proceedings in 2022 [16]. Considering this, this work aimed at providing a second iteration of latter the metabolic model and its respective validation [8]. This model is also to be compared against the subsp. multiplex CFBP 8418 model. Lastly, since there is no effective treatment for infected plants, the GSM model is to be used to investigate potential drugs that could be used as treatment, without affecting olive trees.

## 2. Materials and Methods

### 2.1. Software and Genome Files

A genome-scale information framework (*merlin*) [17] was used to reconstruct the metabolic model. With this software, genome annotation and manual curation can be performed in an easy and intuitive manner, with a variety of features and plugins available to speed up the process. Moreover, COBRApy [18] was used for all simulations associated with the final reconstruction step (model validation). In addition, both COBRApy and Biopython [19] packages were required for the drug targeting pipeline performed in this study.

*merlin* was used to automatically retrieve *X.fastidiosa* subsp. *pauca* De Donno RefSeq assembly genome files, accessible via the NCBI [20] assembly accession number ASM211787v1 [21].

### 2.2. Genome Annotation

The basic local alignment search tool (BLAST) [22] was used to annotate the genome of the bacterium. The similarity searches were performed against UniProtKB/SwissProt and UniProtKB/TrEMBL [**?**], setting an expected value (e-Value) of 1e^-30^ as the threshold.

The *merlin’s* “automatic workflow” tool was used for the annotation, based on phylogenetic proximity to *X.fastidiosa* and the number of records associated with the organism at UniProtKB/SwissProt.

Briefly, the workflow explores the list of retrieved homologous for each candidate gene, comparing it to the defined set of similar organisms. If a match is found, the candidate gene is automatically annotated with the enzyme commission (EC) number [23] and the product name of the closest organism’s homologous gene. The employed automatic workflow is shown in Figure S1, along with the list of selected organisms.

### 2.3. Metabolic Network Reconstruction

#### 2.3.1. Metabolic Data Integration

A draft metabolic network for *X.fastidiosa* was assembled by coupling the information obtained in the genome annotation step with metabolic data retrieved from the Kyoto Encyclopedia of Genes and Genomes (KEGG) database [24]. The EC numbers identified in the genome annotation step were used to select the reactions to be included in the draft network. Reactions involved in generic metabolic pathways, such as DNA methylation and rRNA modification, were ignored and not included in the model as these are unnecessary (modelling-wise) for the reconstruction process. To finalize the initial draft metabolic network, KEGG’s spontaneous reactions were automatically added to the model.

#### 2.3.2. Compartments

The subcellular compartment location of each protein was predicted using PSORTb 3.0 [25]. The resulting file was loaded intomerlin, allowing the automatic integration of compartments into the model’s reactions.

#### 2.3.3. Transport Reactions

Transport reactions were generated using the Transport Systems Tracker (transyt.bio.di.uminho.pt).

#### 2.3.4. Biomass and Energy Requirements

The *Escherichia coli* GSM model (iOJ1366) [26] and *Xanthomonas campestris* large-scale metabolic model [27] were used as templates to infer the macromolecular composition of the biomass equation. This information was complemented with data retrieved from the literature.

The biomass reaction accounts for a total of seven macromolecular entities, namely Deoxyribonucleic acid (DNA), Ribonucleic acid (RNA), Protein, Cofactor, Peptidoglycan, Lipopolysaccharide, and Lipid. Deoxynucleotide, nucleotide, and amino acid contents were determined through *merlin’s* e-Biomass tool [28]. Additionally, this tool formulated a template reaction for the Cofactor macromolecule, which was promptly rearranged considering the metabolic potential of the assembled metabolic network. However, as the composition of the remaining macromolecules cannot be obtained through genomic information, available literature was sought to determine the necessary precursors for the synthesis of peptidoglycans [29], lipopolysaccharides [30,31] and lipids [32]. Finally, as the lipidic portion of the biomass requires fatty acids for its synthesis, whose chain is of variable size, an average fatty acid metabolite was created based on data of multiple strains of *X.fastidiosa* [1] as described elsewhere [33].

Growth and non-growth associated energy requirements were included in the model, as these represent the energy required for several cellular processes, including the assembly of macromolecules and the maintenance of internal homeostasis. As there is a lack of information regarding energy requirements for *X. fastidiosa*, information from the iOJ1366 model was adapted to this work. Therefore, growth-associated energy necessities, represented in the biomass equation, were set as the consumption of 53.5 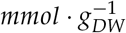 of ATP. Likewise, non-growth associated requirements were defined through an ATP hydrolysis reaction, while setting its lower and upper bounds to 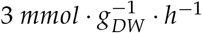.

### 2.4. Manual Curation

To accurately represent an organism’s metabolic behaviour, manual curation is required. Therefore, whenever *in silico* simulations did not corroborate experimental data, manual curation of the GSM model was carried out.

#### 2.4.1. Gap-Filling

Dead-end metabolites and blocked reactions were identified using *merlin*.

Through this phase, an initial assessment of the metabolic potential of *X. fastidiosa* was performed by reviewing organism-specific literature. Specifically, pathways with metabolic gaps were evaluated to a higher degree. Furthermore, whenever a gap was found, supporting evidence at the genome level was sought, in the information obtained from the genome annotation step. If the enzyme that catalyses the missing reaction was indeed encoded in the genome, such reaction, and its associated GPR rule, was added to the model. Thus, multiple missed/misannotations were corrected during this comprehensive analysis.

Moreover, a tool recently added to the *merlin* software, BioISO [34], was also used during this process. BioISO tests the capability of the network towards the synthesis of products of a given reaction. Therefore, this tool is very valuable to test the network potential to produce biomass precursor molecules and can pinpoint potential issues within the network for the users.

### 2.5. Model Validation

The reconstructed GSM model’s performance was evaluated by comparing the predicted results to available information retrieved from the literature. As there is a limited amount of information regarding *X. fastidiosa* metabolism, *X. campestris* data were used whenever necessary. In this work, all *in silico* simulations were made by setting one of the biomass reactions (‘R e-Biomass’ or ‘R Coupled Biomass’) as the objective function and maximizing it using Parsimonious Flux Balance Analysis (pFBA) [35], except otherwise indicated. Finally, multiple metabolic properties were tested (described in the sections below) by restraining the model properly.

#### 2.5.1. Glucose Flux Pattern

Glucose flux pattern through the assembled metabolic network was assessed to allow direct comparison to *X. campestris* experimental data. Therefore, an environmental condition that replicates a minimal medium was formulated, where glucose and ammonia were used as carbon and nitrogen sources, respectively. Additionally, flux through the phosphofructokinase reaction (R00764, EC 2.7.1.90) was restrained to a minimal value (1% of the glucose intake rate), as the pyrophosphate-dependent phosphofructokinase identified in *X. fastidiosa*’s genome displays low processivity and represents a minimal fraction of the total proteome in similar organisms [27,36,37].

A quantitative validation was performed by evaluating the GSM model’s predicted growth and exopolysaccharide production rates. However, as there is limited information related to this topic for *X. fastidiosa*, both data published alongside the large-scale metabolic model of *X. campestris* [27] and reported by Letisse *et al*. [38], were used for validation purposes.

Nonetheless, the environmental conditions had to be properly constrained, identically to previous validation steps, as the data retrieved from *X. campestris* concerns its growth under sucrose, an absent metabolite in the reconstructed GSM model of *X. fastidiosa*. As a result, glucose was used as a carbon source, instead of sucrose, and the carbon intake was normalized according to the number of carbon atoms (an uptake rate of 1.8 *mmol* · *g*^-1^ · *h*^-1^ of sucrose equals an uptake rate of 3.6 *mmol* · *g*^-1^ · *h*^-1^ of glucose).

#### 2.5.2. Comparison with *X. fastidiosa* subsp. multiplex CFBP 8418 GSM Model

For validation purposes, the model developed in this work was directly compared to the GSM model of *X. fastidiosa* subsp. multiplex CFBP 8418 [15]. The first comparison relied on the minimization of glutamine uptake a predefined set of growth, exopolysaccharide production and LesA (a protein secreted by *X. fastidiosa*) production rates. This last entity, a lipase/esterase known to be secreted by *X. fastidiosa* and a key virulence factor during the phytopathogen’s infection process [39], was inserted in the model by creating a biosynthetic reaction that accounted for its amino acid composition.

Distinct carbon sources were tested to examine whether they supported *X. fastidiosa’s* growth. Hence, the environmental conditions replicated a minimal medium, where ammonia, orthophosphate and Fe^2+^ would act as sources of nitrogen, phosphate, and iron, respectively. Additionally, carbon uptake was normalized according to the number of carbon atoms of the carbohydrate. Simulation outputs were then compared to Biolog Phenotype Microarray data, published along with a GSM model of *X. fastidiosa* subsp. multiplex CFBP 8418.

## 3. Results and Discussion

### 3.1. Biomass and Energy Requirements

The macromolecular composition of the *X. fastidiosa’s* model reconstructed in this work, whose entities and respective fractions were inferred from *E. coli* [26] and *X. campestris* [27] models, is compiled in Table 1. A detailed biomass composition is available in Table S1.

**Table 1.** Biomass equation representing the macromolecular composition of *X. fastidiosa* (this work), *E. coli* [26] and *X. campestris* [27] models. The fraction of each macromolecule is represented in grams of molecule per gram of cell dry weight.

| Macromolecule | *E. coli* | *X. campestris* | *X. fastidiosa* |
| --- | --- | --- | --- |
| DNA | 0.031 | 0.040 | 0.040 |
| RNA | 0.205 | 0.160 | 0.160 |
| Protein | 0.550 | 0.490 | 0.490 |
| Lipid | 0.091 | 0.130 | 0.130 |
| Glycogen | 0.025 | -* | - |
| Peptidoglycan | 0.025 | - | 0.0589 |
| Cofactor | 0.039 | - | 0.0706 |
| Lipopolysaccharide | 0.034 | 0.033 | 0.0505 |

\*Although glycogen was not included in the biomass reaction, it's production was constrained to a ratio of 2.21 glycogen/biomass.

Due to the phylogenetic closeness between *X. campestris* and *X. fastidiosa*, most biomass-relative contents were directly retrieved from the former model, whenever possible. Data from *E. coli* was used to determine the remaining macromolecular contents, but the missing portion of *X. campestris* was always considered during the calculations.

Despite the usage of glycogen in iOJ1366 [26] and *X. campestris* [27] models, the macromolecule was not added to the biomass equation of the *X. fastidiosa* model, as the glycogen metabolism seems to be a lost trait in multiple parasitic bacteria, including *X. fastidiosa* [40]. This was further confirmed by the lack of biosynthetic and catabolic enzymes related to glycogen during the genome annotation step. Nevertheless, the glycogen fraction was divided in a 30 – 70% split and allocated into peptidoglycan and lipopolysaccharide, as both these macromolecules contain carbohydrates in their constitution. DNA, RNA and Protein compositions, in terms of precursors, were determined using the e-Biomass tool [28], which is embedded in*merlin*.

Peptidoglycan production was represented by creating a reaction that accounted for all of its precursors. As *X. fastidiosa*’s peptidoglycan data is still lacking, information from *E. coli* [29] was retrieved to formulate peptidoglycan biosynthesis, which is composed of two monosaccharide units and five amino acids. One of these amino acids is D-Glutamate, which was promptly replaced by the enantiomer L-Glutamate as *X.fastidiosa’s* metabolic network could not support its biosynthesis.

The lipidic portion of the biomass, which comprises membrane phospholipids, was adapted from data of *X. campestris* B-24 [32], as no information was available for *X. fastidiosa*. Phosphatidylinositol and lysophosphatidylethanolamine, compounds measured in the previous experiment, were ignored since these were not found in the assembled *X. fastidiosa*’s metabolic network. Therefore, these lipidic contents were distributed among the other phospholipids. LPS, a constituent of gram-negative bacterial outer membranes, is composed of three moieties: Lipid A, core oligosaccharide and O-antigen. While the Lipid A and the inner portion of the core oligosaccharide are conserved in different bacterial species, both the core oligosaccharide’s outer section and the O-antigen, which is an important virulence determinant, are structurally variable [30,31]. These structural changes were inferred from data published by [29] and an LPS synthesis reaction specific for *X. fastidiosa* was assembled.

The remaining biomass portion was associated with the e-Cofactor macromolecule, which is first determined using the e-Biomass tool [28]. However, a rearrangement was performed according to the metabolic potential of the network. As no information regarding the amount of each molecule in the cofactor pool is available, it was assumed that each cofactor would have the same weight (in grams) in the final biomass formulation.

The inclusion of energy requirements, which include both cell growth and maintenance-related energy, completed the *X. fastidiosa’s* biomass formulation. As stated previously, energy necessities were retrieved from the iOJ1366 *E. coli* model [26].

Due to the possibility of exopolysaccharide biosynthesis during the life cycle of *X. fastidiosa*, a reaction that couples biomass and exopolysaccharide production was inserted in the model (“R_Coupled_Biomass”), as these entities would compete for the available carbon source. This reaction forces the synthesis of biomass and fastidian gum (X. *fastidiosa’s* exopolysaccharide) as follows: 0.203 Biomass+0.797 Fastidian gum=>1.0 Coupled Biomass.

The stoichiometries were inferred from published data for *X. campestris* [41], and employed in the large-scale metabolic model of *X. campestris* [27].

### 3.2. Model Validation

#### 3.2.1. Glucose Flux Pattern

Several xanthomonads use the Entner-Doudoroff pathway (EMP) as the main catabolic route for glucose [42], including *X. campestris* [37]. Although this was not demonstrated for *X. fastidiosa* yet, due to the genetic and metabolic similarities among these organisms, the EMP has been proposed to be the main glucose degradation pathway for *X. fastidiosa* [43].

By assuming the maximisation of biomass production, the model predicts 88.6% and 8.6% of the glucose flux towards the Entner-Doudoroff and the non-oxidative branch of the pentose phosphate pathways, respectively (Figure 1), which is in the range of the experimental data available for other organisms: 81 – 93% and 7 – 19% [42]. A more detailed analysis of the glucose flux pattern from this model was already published previously [16].

**Figure 1.**
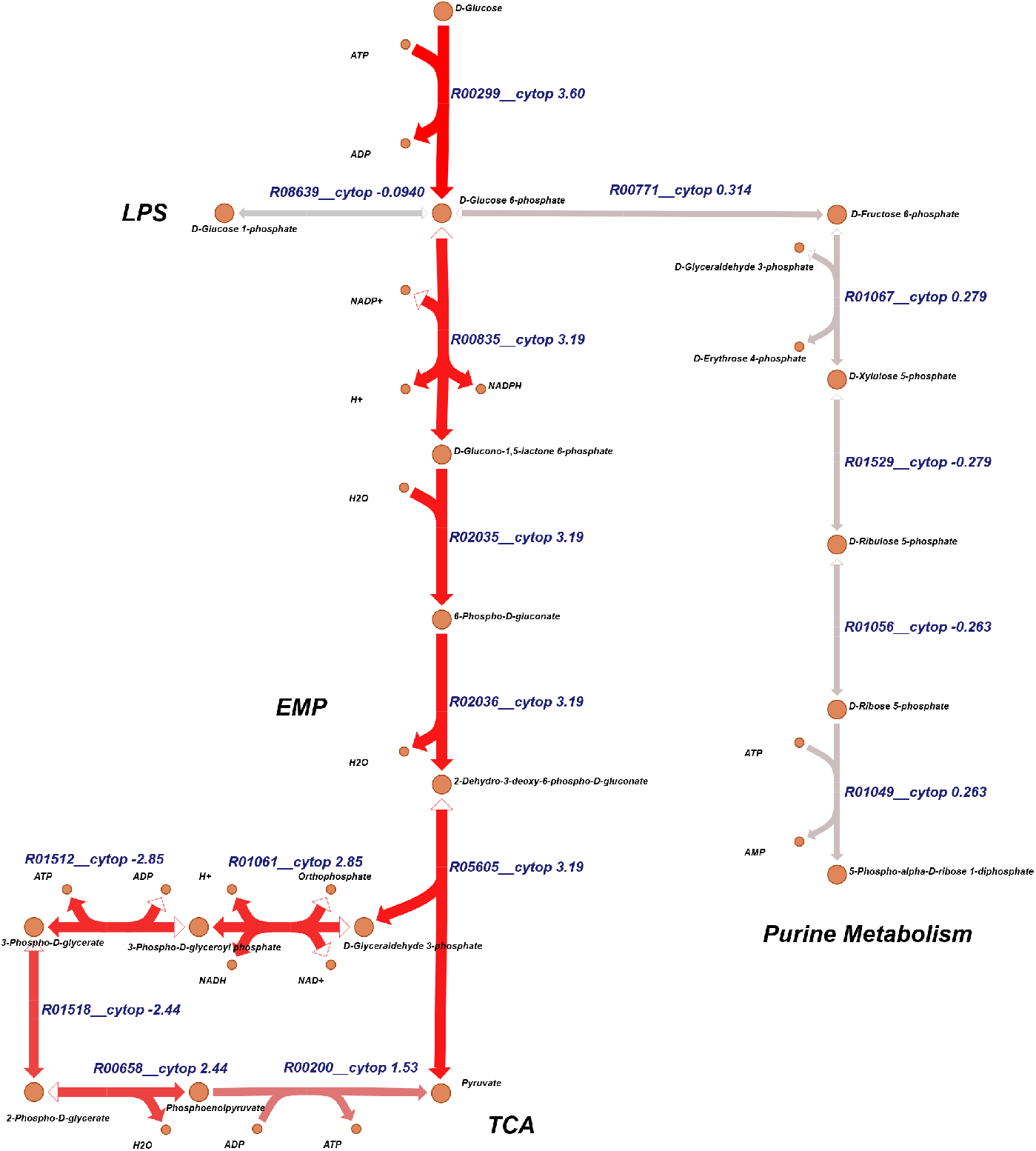
Glucose catabolism obtained from a pFBA simulation.

#### 3.2.2. Specific growth rate and exopolysaccharide production

Specific growth and exopolysaccharides production rates were directly compared with *in silico* simulations of the *X. campestris* large scale metabolic model [27] and experimental data of the aforementioned species [38]. *X.fastidiosa* subsp. *pauca* De Donno GSM model was properly constrained to be possible to perform this direct comparison. Therefore, glucose was used for the *in silico* simulations of the model developed in this work.

As displayed in table 2, the model *X.fastidiosa* predicted a growth rate value of 0.11 *h*^−1^ and a specific production rate of exopolysaccharide of 0.44 *mmol* · *g*^−1^ · *h*^−1^. The results are similar to the experimental data [38] and agree with the known fact that *X. fastidiosa* displays slower growth and exopolysaccharide production rates in comparison to *X. campestris*.

**Table 2.**
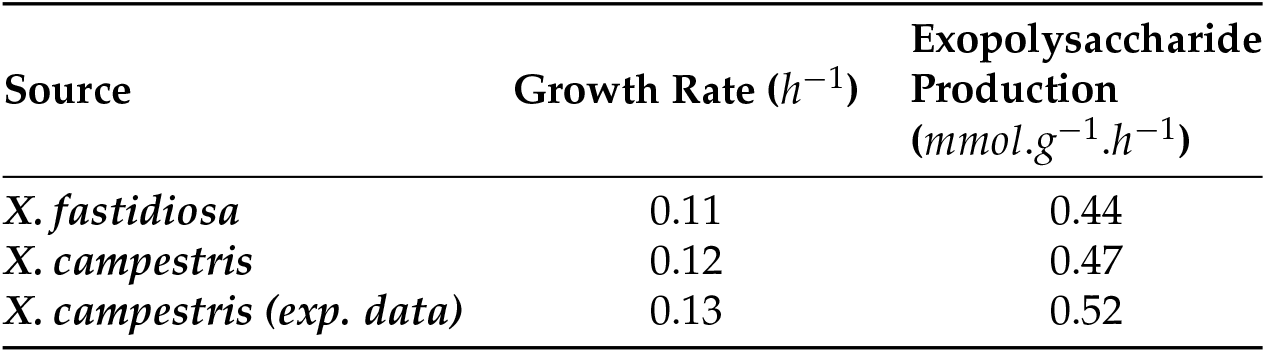
*In silico* predictions of growth and exopolysaccharide production rates for *X.fastidiosa* iMS509, and *X. campestris* [27] metabolic models, in comparison to experimental data specific for *X. campestris* [38].

The other end-products of *X. fastidiosa* subsp. *pauca* De Donno GSM model *in silico* simulations were assessed. As expected, the major outputs are water and carbon dioxide, which are by-products of the respiration metabolism. On the other hand, xanthomonadin was predicted to be produced at a small rate. Note that these metabolites were inferred to be produced by the organism after an analysis of the metabolic capabilities of the assembled network. Firstly, xanthomonadins are yellow pigments with photobiological damage protection properties produced by *Xanthomonas spp*. [44]. Since *X.fastidiosa* lacks a transaldolase (EC 2.2.1.2), D-Erythrose-4-phosphate can not be completely recycled by the organism; hence, xanthomonadins production was inferred. According to the metabolic model, D-Erythrose-4-phosphate could only be metabolised in the shikimate pathway and, as a result, dead-end metabolites of this pathway were thoroughly analysed. Evidence found shows that chorismate was linked to the synthesis of both ubiquinone and xanthomonadin through a bifunctional chorismatase [45] found in *X. fastidiosa*’s genome. Therefore, we assumed the production of xanthomonadin and added the necessary reactions.

### 3.3. Model Summary

The GSM model reconstructed in this work, comprises a total of 509 genes, 1234 metabolites and 1138 reactions, whose distribution can be seen in Figure 2. This model also includes 977 reactions with GPR associations, 248 transport reactions, and 90 exchange reactions. The iMS509 model is available in the BioModels database [46] (MODEL2212050001) in Systems Biology Markup Language (SBML) version 3 format [47].

**Figure 2.**
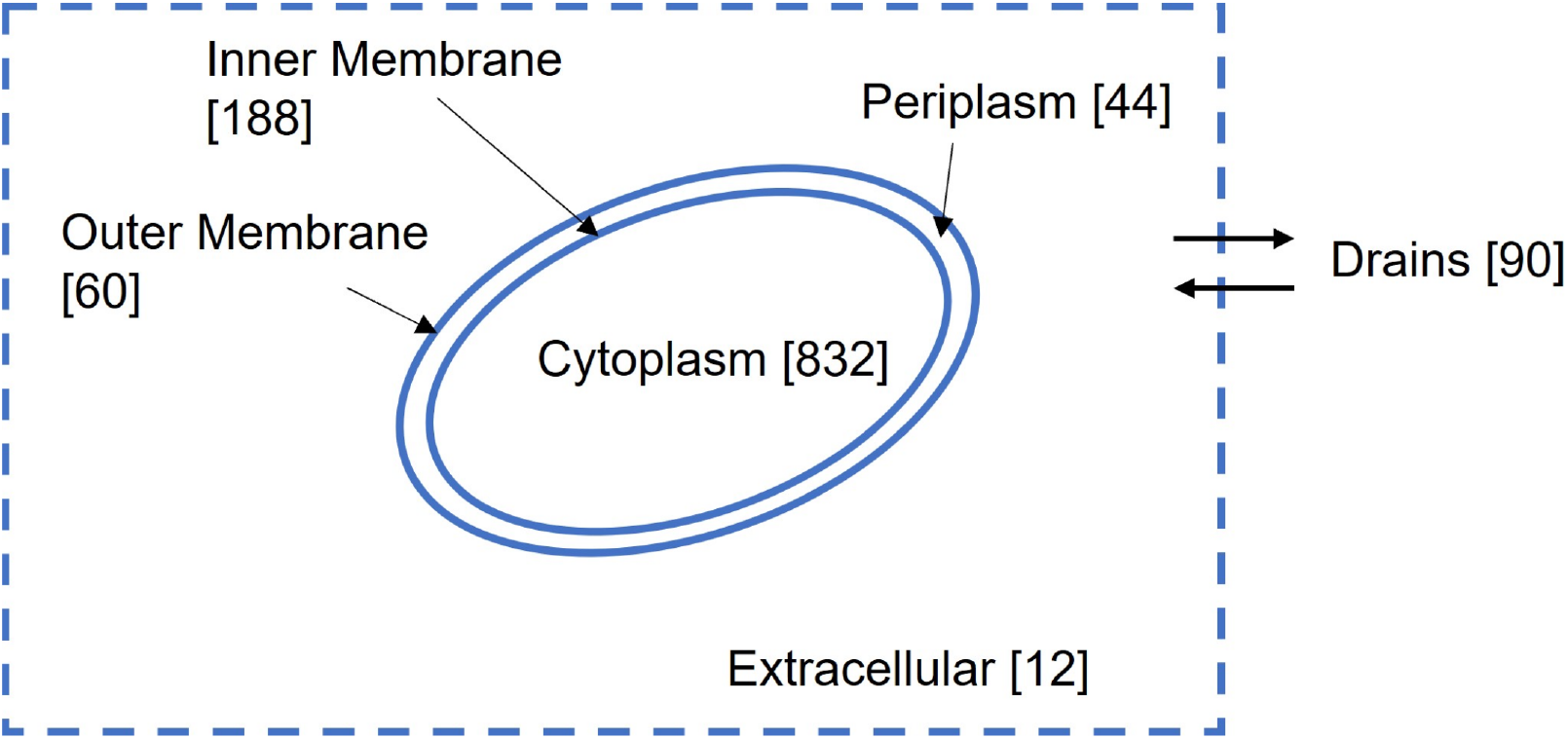
Summary of the distribution of reactions among the different compartments in the iMS498 model, which accounts for a total of 1164 metabolic reactions and 90 drains.

### 3.4. Comparison with X.fastidiosa multiplex CFBP 8418 GSM model

Recently, a GSM model of *X. fastidiosa* multiplex CFBP 8418 was published [15]. Therefore, the two metabolic models were compared to assess metabolic differences between the two strains. A topological comparison between the models is displayed in Table 3, and as it can be seen, the models show a small difference in the number of genes, metabolites, and reactions.

**Table 3.** Summary of *X. fastidiosa* subsp. *pauca* De Donno (this work) and *X. fastidiosa* multiplex CFBP 8414 [15] models, describing the number of genes, reactions, and metabolites.

|  | *X. fastidiosa*<br>*pauca* De Donno | *X. fastidiosa*<br>multiplex CFBP 8418 |
| --- | --- | --- |
| **Genes** | 509 | 572 |
| **Metabolites** | 1234 | 1107 |
| **Reactions** | 1138 | 1158 |

Subsequently, a minimisation of L-glutamine uptake, while restraining the growth rate, EPS and LesA production to a specific output, was performed and compared to the published data. As displayed in Table 4, the uptake of L-glutamine and other metabolites was identical, and the discrepancies can be explained by the intrinsic details of the biomass assembly reaction of each model. Additionally, the composition of the LesA protein is different in these two models; however, the prediction results do not show any significant difference between both models.

**Table 4.**
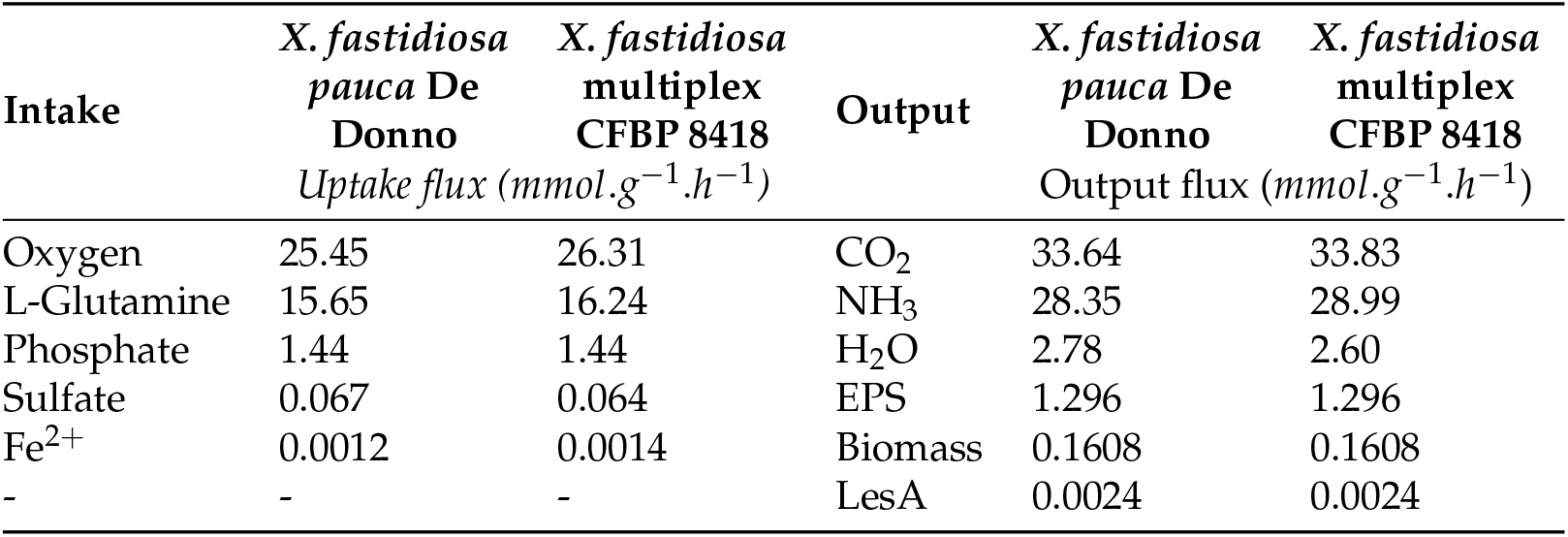
Simulation performance between the two *X. fastidiosa’s* GSM models, in minimisation of L-glutamine intake situation.

| Intake | *X. fastidiosa*<br>*pauca* De<br>Donno<br>Uptake flux ($\text{mmol.g}^{-1}.\text{h}^{-1}$) | *X. fastidiosa*<br>multiplex<br>CFBP 8418 | Output | *X. fastidiosa*<br>*pauca* De<br>Donno<br>Output flux ($\text{mmol.g}^{-1}.\text{h}^{-1}$) | *X. fastidiosa*<br>multiplex<br>CFBP 8418 |
| --- | --- | --- | --- | --- | --- |
| Oxygen | 25.45 | 26.31 | CO<sub>2</sub> | 33.64 | 33.83 |
| L-Glutamine | 15.65 | 16.24 | NH<sub>3</sub> | 28.35 | 28.99 |
| Phosphate | 1.44 | 1.44 | H<sub>2</sub>O | 2.78 | 2.60 |
| Sulfate | 0.067 | 0.064 | EPS | 1.296 | 1.296 |
| Fe<sup>2+</sup> | 0.0012 | 0.0014 | Biomass | 0.1608 | 0.1608 |
| - | - | - | LesA | 0.0024 | 0.0024 |

A comparison to evaluate the usage of different carbon sources by *X. fastidiosa* was then performed. The results revealed different behaviours between both models for a subset of the tested carbon sources (Figure 3). While *X. fastidiosa* subsp. *pauca* De Donno GSM model only predicts growth for 14 out of the 24 tested carbon sources, the *X. fastidiosa* multiplex CFPB 8414 GSM model simulates growth in all of them. *In silico* simulations of the iMS509 model do not indicate growth for the substrates L-Proline, L-Arginine, L-Histidine, L-Ornithine, GABA, Acetate, D-Galactose, D-Xylose, and myo-Inositol, due to the lack of genomic evidence concerning their catabolic pathways. On the other hand, the model published by Gerlin and colleagues [15] includes the necessary reactions, although without any genetic association, which may be valid for *X. fastidiosa* multiplex CFPB 8418 according to the data provided by Biolog Phenotype Microarrays, published alongside the model. Nevertheless, the same behaviour assumption could not be used for the strain used in this work. Not only due to the lack of genomic evidence, but also because other strains do not show growth in media supplemented with L-Histidine, L-Arginine and L-Lysine [48]. Furthermore, it is common for different strains of the same organism to display distinct metabolic behaviours, as shown in *Streptococcus pneumoniae* [33], which can be caused by both genetic diversity and gene regulation. Hence, we assumed that *X. fastidiosa* subsp. *pauca* De Donno was unable to degrade these compounds and the model was reconstructed accordingly.

**Figure 3.**
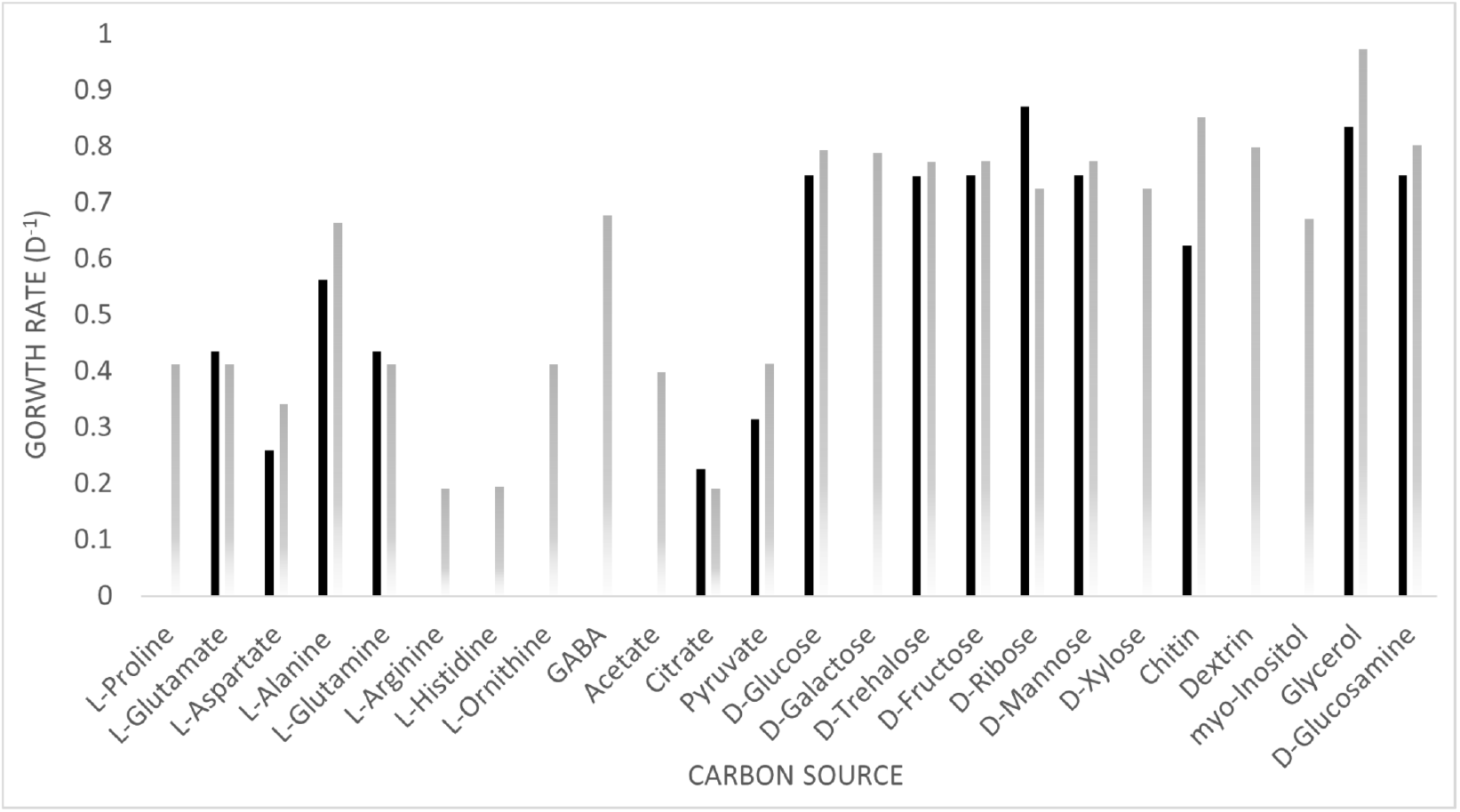
Comparison of *X.fastidiosa*’s GSM models growth rate (*d*^−1^) predictions under different carbon sources. The lack of growth prediction in some carbon sources by the *X. fastidiosa pauca* De Donno GSM model is justified by the lack of genomic evidence for enzymes related to the catabolic pathways of the metabolites, while the other model included these catabolic reactions without any gene association.

As stated previously, the presence of a pyrophosphate-dependent phosphofructokinase (EC 2.7.1.90) was identified, allowing the usage of gluconeogenesis by the phytopathogenic organism. However, only the model developed in this work contains this gene-enzyme-reaction association, while in *X. fastidiosa* multiplex CFBP 8418 it was assumed the utilization of this pathway, through the more typical enzyme fructose-1,6-bisphosphate (EC 3.1.3.11) although without any genetic evidence/association to this enzyme.

## 4. Conclusions

The output of this work was the reconstruction of a GSM model for *X. fastidiosa* subsp. *pauca* De Donno, named iMS509. This work was a second iteration of a previouly developed model named iMS508 [16] The model was reconstructed using *merlin*, using an automatic genome annotation approach. Biological databases were also considered during the process of retrieving relevant information. The metabolic model has 509 genes, 1234 metabolites, 1138 reactions, and 3 compartments. The model was successfully validated accordingly to the glucose flux pattern. Moreover, it was able to simulate aerobic growth of the bacterium and accurately predict production rates of fastidian gum and the lesA protein.

## Supplementary Materials

Figure S1: Automatic workflow employed; Figure S2: murA aminoacid sequence; Table S1: Detailed biomass composition of *X. fastidiosa pauca* De Donno; Table S2: Genome annotation results; Table S3: Complete list of drugs known to inhibit the found targets.

## Author Contributions

Conceptualization: AO, EC, MS, CF, and OD; Data curation: AO, EC, and MS; Formal analysis: AO, EC, and MS; Methodology: AO, EC, MS, CF, and OD; Supervision: OD and CF; Validation: AO, EC; Writing - original draft: AO and EC; Writing - review & editing: OD and CF

## Data Availability Statement

The model presented in this work can be found in BioModels with the identifier MODEL2212050001.

## Acknowledgments

We would like to acknowledge the Portuguese Foundation for Science and Technology for supporting this work.

## Abbreviations

The following abbreviations are used in this manuscript:

BLAST: Basic Local Alignment Search Tool
EMP: Entner-Doudoroff pathway
GSM: Genome-scale metabolic
I2P2: Impact Indicator for Priority Pests
*merlin*: Metabolic Models Reconstruction Using Genome-Scale Information
PBP: Penicillin-Binding Protein
ROS: Reactive Oxygen Species
SBML: Systems Biology Markup Language

## References

1. Wells, J.M.; Raju, B.C.; Hung, H.Y.; Weisburg, W.G.; Mandelco-Paul, L.; Brenner, D.J. *Xylella fastidiosa* gen. nov., sp. nov: Gram-Negative, Xylem-Limited, Fastidious Plant Bacteria Related to *Xanthomonas* spp. International Journal of Systematic Bacteriology 1987, 37, 136–143. https://doi.org/10.1099/00207713-37-2-136.

2. Chatterjee, S.; Almeida, R.P.P.; Lindow, S. Living in two Worlds: The Plant and Insect Lifestyles of *Xylella fastidiosa*. Annual Review of Phytopathology 2008, 46, 243–271. https://doi.org/10.1146/annurev.phyto.45.062806.094342.

3. Bradbury, J. Guide to Plant Pathogenic Bacteria; CAB International Mycological Institute, 1986.

4. (EFSA), E.F.S.A. Update of the *Xylella* spp. host plant database, 2018. https://doi.org/10.2903/j.efsa.2018.5408.

5. Nunney, L.; Vickerman, D.B.; Bromley, R.E.; Russell, S.A.; Hartman, J.R.; Morano, L.D.; Stouthamer, R. Recent Evolutionary Radiation and Host Plant Specialization in the *Xylella fastidiosa* Subspecies Native to the United States. Applied and Environmental Microbiology 2013, 79, 2189–2200. https://doi.org/10.1128/AEM.03208-12.

6. Baldi, P.; Porta, N.L. *Xylella fastidiosa*: Host Range and Advance in Molecular Identification Techniques. Frontiers in Plant Science 2017, 8, 1–22. https://doi.org/10.3389/fpls.2017.00944.

7. Elbeaino, T.; Valentini, F.; Abou Kubaa, R.; Moubarak, P.; Yaseen, T.; Digiaro, M. Multilocus sequence typing of *Xylella fastidiosa;* isolated from olive affected by “olive quick decline syndrome” in Italy. Phytopathologia Mediterranea 2014, 53, 533–542. https://doi.org/10.14601/Phytopathol_Mediterr-15000.

8. Saponari, M.; Giampetruzzi, A.; Loconsole, G.; Boscia, D.; Saldarelli, P. *Xylella fastidiosa* in olive in Apulia: where we stand. Phytopathology 2018, pp. 1–43. https://doi.org/10.1094/PHYTO-08-18-0319-FI.

9. Schneider, K.; van der Werf, W.; Cendoya, M.; Mourits, M.; Navas-Cortés, J.A.; Vicent, A.; Lansink, A.O. Impact of *Xylella fastidiosa* subspecies *pauca* in European olives. Proceedings of the National Academy of Sciences 2020, 117, 9250–9259. https://doi.org/10.1073/pnas.1912206117.

10. Vos, S.; Camilleri, M.; Diakaki, M.; Lázaro, E.; Parnell, S.; Schrader, G.; Vicent, A. Pest survey card on Xylella fastidiosa. EFSA Supporting Publications 2019. https://doi.org/10.2903/sp.efsa.2019.EN-1667.

11. Palsson, B. Systems Biology - Properties of Reconstructed Networks; 2006.

12. Dias, O.; Rocha, I., Systems biology in fungi. In Molecular Biology of Food and Water Borne Mycotoxigenic and Mycotic Fungi; CRC Press Boca Raton, 2015; pp. 69–92.

13. Presta, L.; Bosi, E.; Mansouri, L.; Dijkshoorn, L.; Fani, R.; Fondi, M. Constraint-based modeling identifies new putative targets to fight colistin-resistant *A. baumannii* infections. Scientific Reports 2017, 7, 3706. https://doi.org/10.1038/s41598-017-03416-2.

14. Cesur, M.F.; Siraj, B.; Uddin, R.; Durmuş, S.; Çakir, T. Network-Based Metabolism-Centered Screening of Potential Drug Targets in *Klebsiella pneumoniae* at Genome Scale. Frontiers in cellular and infection microbiology 2019, 9, 447. https://doi.org/10.3389/fcimb.2019.00447.

15. Gerlin, L.; Cottret, L.; Cesbron, S.; Taghouti, G.; Jacques, M.A.; Genin, S.; Baroukh, C. Genome-Scale Investigation of the Metabolic Determinants Generating Bacterial Fastidious Growth. mSystems 2020, 5, 1–15. https://doi.org/10.1128/mSystems.00698-19.

16. Oliveira, A.; Cunha, E.; Silva, M.; Faria, C.; Dias, O. Exploring Xylella fastidiosa’s Metabolic Traits Using a GSM Model of the Phytopathogenic Bacterium. In Proceedings of the Practical Applications of Computational Biology and Bioinformatics, 16th International Conference (PACBB 2022); Fdez-Riverola, F.; Rocha, M.; Mohamad, M.S.; Caraiman, S.; Gil-González, A.B., Eds.; Springer International Publishing: Cham, 2023; pp. 79–88.

17. Capela, J.; Lagoa, D.; Rodrigues, R.; Cunha, E.; Cruz, F.; Barbosa, A.; Bastos, J.; Lima, D.; Ferreira, E.C.; Rocha, M.; et al. merlin, an improved framework for the reconstruction of high-quality genome-scale metabolic models. Nucleic Acids Research 2022, 50, 6052–6066. https://doi.org/10.1093/nar/gkac459.

18. Ebrahim, A.; Lerman, J.A.; Palsson, B.O.; Hyduke, D.R. COBRApy: constraints-based reconstruction and analysis for python. BMC systems biology 2013, 7, 1–6. https://doi.org/10.1186/1752-0509-7-74.

19. Cock, P.J.A.; Antao, T.; Chang, J.T.; Chapman, B.A.; Cox, C.J.; Dalke, A.; Friedberg, I.; Hamelryck, T.; Kauff, F.; Wilczynski, B.; et al. Biopython: freely available Python tools for computational molecular biology and bioinformatics. Bioinformatics 2009, 25, 1422–1423. https://doi.org/10.1093/bioinformatics/btp163.

20. Sayers, E.W.; Bolton, E.E.; Brister, J.R.; Canese, K.; Chan, J.; Comeau, D.; Connor, R.; Funk, K.; Kelly, C.; Kim, S.; et al. Database resources of the national center for biotechnology information. Nucleic Acids Research 2022, 50, D20–D26. https://doi.org/10.1093/NAR/GKAB1112.

21. Giampetruzzi, A.; Saponari, M.; Almeida, R.P.P.; Essakhi, S.; Boscia, D.; Loconsole, G.; Saldarelli, P. Complete Genome Sequence of the Olive-Infecting Strain *Xylella fastidiosa* subsp. *pauca* De Donno. Genome Announcements 2017, 5, 5–6. https://doi.org/10.1128/genomeA.00569-17.

22. Altschul, S.F.; Gish, W.; Miller, W.; Myers, E.W.; Lipman, D.J. Basic local alignment search tool. Journal of Molecular Biology 1990, 215, 403–410. https://doi.org/10.1016/S0022-2836(05)80360-2.

23. Enzyme nomenclature 1992: recommendations of the Nomenclature Committee of the International Union of Biochemistry and Molecular Biology on the nomenclature and classification of enzymes: International Union of Biochemistry and Molecular Biology. Nomenclature Committee, author: Free Download, Borrow, and Streaming: Internet Archive.

24. Kanehisa, M.; Furumichi, M.; Tanabe, M.; Sato, Y.; Morishima, K. KEGG: new perspectives on genomes, pathways, diseases and drugs. Nucleic Acids Research 2016, 45, D353–D361. https://doi.org/10.1093/nar/gkw1092.

25. Yu, N.Y.; Wagner, J.R.; Laird, M.R.; Melli, G.; Rey, S.; Lo, R.; Dao, P.; Sahinalp, S.C.; Ester, M.; Foster, L.J.; et al. PSORTb 3.0: improved protein subcellular localization prediction with refined localization subcategories and predictive capabilities for all prokaryotes. Bioinformatics 2010, 26, 1608–1615. https://doi.org/10.1093/bioinformatics/btq249.

26. Orth, J.D.; Conrad, T.M.; Na, J.; Lerman, J.A.; Nam, H.; Feist, A.M.; Palsson, B.Ø. A comprehensive genome-scale reconstruction of *Escherichia coli* metabolism. Molecular Systems Biology 2011, 7, 1–9. https://doi.org/10.1038/msb.2011.65.

27. Schatschneider, S.; Persicke, M.; Watt, S.A.; Hublik, G.; Pühler, A.; Niehaus, K.; Vorhölter, F.J. Establishment, *in silico* analysis, and experimental verification of a large-scale metabolic network of the xanthan producing *Xanthomonas campestris* pv. campestris strain B100. Journal of Biotechnology 2013, 167, 123–134. https://doi.org/10.1016/j.jbiotec.2013.01.023.

28. Santos, S.; Rocha, I. Estimation of biomass composition from genomic and transcriptomic information. Journal of Integrative Bioinformatics 2016, 13, 1–14. https://doi.org/10.1515/jib-2016-285.

29. Vollmer, W.; Holtje, J.V. The Architecture of the Murein (Peptidoglycan) in Gram-Negative Bacteria: Vertical Scaffold or Horizontal Layer (s)? Journal of Bacteriology 2004, 186, 5978–5987. https://doi.org/10.1128/JB.186.18.5978.

30. Wang, X.; Quinn, P.J. Lipopolysaccharide: Biosynthetic pathway and structure modification. Progress in Lipid Research 2010, 49, 97–107. https://doi.org/10.1016/j.plipres.2009.06.002.

31. Whitfield, C. Biosynthesis of lipopolysaccharide O antigens. Trends in Microbiology 1995, 3, 178–185. https://doi.org/10.1016/s0966-842x(00)88917-9.

32. Dianese, J.C.; Schaad, N.W. Isolation and Characterization of Inner and Outer Membranes of *Xanthomonas campestris* pv. campestris. Phytopathology 1982, 72, 1284–1289. https://doi.org/10.1128/jb.176.11.3354-3359.1994.

33. Dias, O.; Saraiva, J.; Faria, C.; Ramirez, M.; Pinto, F.; Rocha, I. iDS372, a Phenotypically Reconciled Model for the Metabolism of Streptococcus pneumoniae Strain R6. Frontiers in Microbiology 2019, 10. https://doi.org/10.3389/fmicb.2019.01283.

34. Cruz, F.; Capela, J.; Ferreira, E.C.; Rocha, M.; Dias, O. BioISO: an objective-oriented application for assisting the curation of genome-scale metabolic models. bioRxiv 2021, [https://www.biorxiv.org/content/early/2021/03/12/2021.03.07.434259.full.pdf]. https://doi.org/10.1101/2021.03.07.434259.

35. Lewis, N.E.; Hixson, K.K.; Conrad, T.M.; Lerman, J.A.; Charusanti, P.; Polpitiya, A.D.; Adkins, J.N.; Schramm, G.; Purvine, S.O.; Lopez-Ferrer, D.; et al. Omic data from evolved *E. coli* are consistent with computed optimal growth from genome-scale models. Molecular systems biology 2010, 6, 390. https://doi.org/10.1038/msb.2010.47.

36. Frese, M.; Schatschneider, S.; Voss, J.; Vorhölter, F.J.; Niehaus, K. Characterization of the pyrophosphate-dependent 6-phosphofructokinase from *Xanthomonas campestris* pv. campestris. Archives of Biochemistry and Biophysics 2014, 546, 53–63. https://doi.org/10.1016/j.abb.2014.01.023.

37. Schatschneider, S.; Huber, C.; Neuweger, H.; Watt, T.F.; Pühler, A.; Eisenreich, W.; Wittmann, C.; Niehaus, K.; Vorhölter, F.J. Metabolic flux pattern of glucose utilization by *Xanthomonas campestris* pv. campestris: prevalent role of the Entner – Doudoroff pathway and minor fluxes through the pentose phosphate pathway and glycolysis. Molecular BioSystems 2014, 10, 2663–2676. https://doi.org/10.1039/C4MB00198B.

38. Letisse, F.; Chevallereau, P.; Simon, J.L.; Lindley, N.D. Kinetic analysis of growth and xanthan gum production with *Xanthomonas campestris* on sucrose, using sequentially consumed nitrogen sources. Applied Microbiology and Biotechnology 2001, 55, 417–422. https://doi.org/10.1007/s002530000580.

39. Nascimento, R.; Gouran, H.; Chakraborty, S.; Gillespie, H.W.; Almeida-Souza, H.O.; Tu, A.; Rao, B.J.; Feldstein, P.A.; Bruening, G.; Goulart, L.R.; et al. The Type II Secreted Lipase/Esterase LesA is a Key Virulence Factor Required for *Xylella fastidiosa* Pathogenesis in Grapevines. Scientific Reports 2016, 6, 1–17. https://doi.org/10.1038/srep18598.

40. Henrissat, B.; Deleury, E.; Coutinho, P.M. Glycogen metabolism loss: a common marker of parasitic behaviour in bacteria? Trends in Genetics 2002, 18, 437–440. https://doi.org/10.1016/S0168-9525(02)02734-8.

41. Pielken, P.; Schimz, K.L.; Eggeling, L.; Sahm, H. Glucose metabolism in *Xanthomonas campestris* and influence of methionine on the carbon flow. Canadian Journal of Microbiology 1988, 34, 1333–1337. https://doi.org/10.1139/m88-234.

42. Zagallo, A.C.; Wang, C.H. Comparative Glucose Catabolism of *Xanthomonas* Species. Journal of Bacteriology 1967, 93, 970–975. https://doi.org/10.1128/jb.93.3.970-975.1967.

43. Facincani, A.P.; Ferro, J.A.; Jr., J.M.P.; Jr., H.A.P.; de Macedo Lemos, E.G.; do Prado, A.L.; Ferro, M.I.T. Carbohydrate metabolism of *Xylella fastidiosa:* Detection of glycolytic and pentose phosphate pathway enzymes and cloning and expression of the enolase gene. Genetics and Molecular Biology 2003, 26, 203–211. https://doi.org/10.1590/S1415-47572003000200015.

44. Poplawsky, A.R.; Urban, S.C.; Chun, W. Biological Role of Xanthomonadin Pigments in *Xanthomonas campestris* pv. Campestris. Applied and Environmental Microbiology 2000, 66, 5123–5127. https://doi.org/10.1128/AEM.66.12.5123-5127.2000.

45. Zhou, L.; Wang, J.Y.; Wang, J.; Poplawsky, A.; Lin, S.; Zhu, B.; Chang, C.; Zhou, T.; Zhang, L.H.; He, Y.W. The diffusible factor synthase XanB2 is a bifunctional chorismatase that links the shikimate pathway to ubiquinone and xanthomonadins biosynthetic pathways. Molecular Microbiology 2013, 87, 80–93. https://doi.org/10.1111/mmi.12084.

46. Malik-Sheriff, R.S.; Glont, M.; Nguyen, T.V.N.; Tiwari, K.; Roberts, M.G.; Xavier, A.; Vu, M.T.; Men, J.; Maire, M.; Kananathan, S.; et al. BioModels — 15 years of sharing computational models in life science. Nucleic Acids Research 2020, 48, D407–D415. https://doi.org/10.1093/nar/gkz1055.

47. Hucka, M.; Bergmann, F.T.; Dräger, A.; Hoops, S.; Keating, S.M.; Novère, N.L.; Myers, C.J.; Olivier, B.G.; Sahle, S.; Schaff, J.C.; et al. The Systems Biology Markup Language (SBML): Language Specification for Level 3 Version 2 Core. Journal of Integrative Bioinformatics 2018, 15, 20170081. doi: https://doi.org/10.1515/jib-2017-0081.

48. Chang, C.J.; Donaldson, R.C. Nutritional requirements of *Xylella fastidiosa*, which causes Pierce’s disease in grapes. Canadian Journal of Microbiology 2000, 46, 291–293. https://doi.org/10.1139/w99-141.

