## Supplementary material for "An updated genome-scale model for *Xylella fastidiosa* subsp. *pauca* De Donno": Figure S1

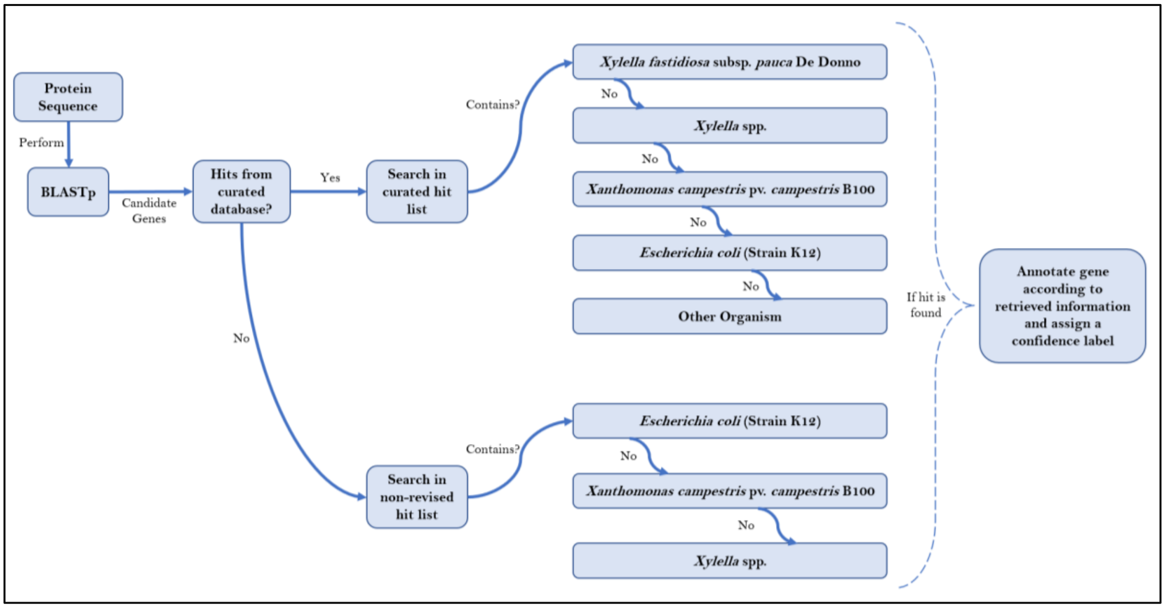


Figure S1 - Annotation workflow followed throughout the genome annotation step. Firstly, all coding sequences found in the organisms' genome is submitted to a BLASTp. Subsequently, only candidate genes go through the automatic annotation process, where the retrieved hit list is searched in terms of source organism and compared to a set of defined organisms. Upon detection of a match, the gene is annotated according to the data retrieved from the hit.
