## Supplementary material for "An updated genome-scale model for *Xylella fastidiosa* subsp. *pauca* De Donno": Figure S2

```
>WP_010893916.1  UDP-N-acetylglucosamine  1-carboxyvinyltransferase  [Xylella
fastidiosa]
MSKIVVAGGTPLCGDVRISGAKNAVLPILSATLLADAPVEISNVPYLHDVITMINLLRELGAGVTMNEGI
EAKGRSITVDPRWVRQHMVPYDLVKTMRASVLLLGPLLACYGAAEVALPGGCAIGSRPVDQHIRGLQSLG
AEITVENGYIKASVSQGRLKGGRFVFDVVSVTGTENLLMAAAVAQGTSVIENAAMEPEVVDLAECLIALG
ARIEGAGTPRIVVEGVERLKGGQYAVLPDRIETGTFLVATAMTGGRISMQQVRPQTLDAVLGKLTEAGAC
IEIGEDSIRLDMQGRRPCSVNLTTAPYPGFPTDMQAQFMALNCVAEGVGVIKETIFENRFMHVDELLRLG
AKIQIEGHTAIVQGVERLSGAPVMATDLRASASLILAGLVAEGETIIDRIYHLDRGYENIEGKLGALGAS
IRRMT
```

Figure S2 – Protein sequence of UDP-N-acetylglucosamine 1-carboxyvinyltransferase (murA) gene, highlighting the cysteine residue in the active site.
